# A cleavable signal peptide controls the topology and Golgi targeting of the membrane protein TMEM165

**DOI:** 10.64898/2026.08.19.745746

**Authors:** Marie-Odile Velings, Romain Simar, Andréa Bleret, Virginie Tevel, Marielle Boonen, Pierre Morsomme

## Abstract

TMEM165 is a Golgi-resident multi-pass membrane protein involved in divalent cation homeostasis and associated with congenital disorders of glycosylation, yet its N-terminal biogenesis has remained unresolved. Here, we demonstrate that TMEM165 contains a functional cleavable signal peptide required for correct Golgi targeting and membrane topology. Loss of this signal peptide causes protein mislocalization, and altered topology with N-terminal cytosolic exposure, whereas extended N-terminal deletion restores both Golgi localization and overall membrane topology, consistent with insertion mediated by the first transmembrane domain as commonly described for multi-pass membrane proteins. Importantly, this N-terminally truncated form remains responsive to manganese-induced degradation and partially restores glycosylation defects associated with TMEM165 deficiency, indicating that the extended N-terminal region is dispensable for core TMEM165 function. Together, these findings identify the signal peptide as a key determinant of TMEM165 biogenesis and suggest that its conservation may contribute not only to membrane targeting, but also to maintaining the proper luminal environment of the N-terminus during early biogenesis.

**Summary:** TMEM165, a Golgi cation transporter linked to a glycosylation disorder, requires its N-terminal signal peptide for correct topology and Golgi targeting; without it, the protein flips orientation and mislocalizes to the ER. The extended N-terminal region beyond the signal peptide, however, is dispensable for TMEM165’s function.

## Introduction

Membrane proteins constitute a major and functionally diverse class of cellular proteins. It has been estimated that approximately 25% of protein-coding genes in all organisms encode membrane-spanning proteins, which play essential roles in ion and nutrient transport, cell signaling, pathogen– host interactions, defense mechanisms and cell adhesion [1]. In eukaryotic cells, the vast majority of these proteins are inserted into the membrane of the endoplasmic reticulum (ER), where they undergo folding, maturation and assembly prior to trafficking to their final cellular destinations [2].

The biogenesis of integral membrane proteins can be conceptually divided into four tightly interconnected steps: targeting, insertion, folding and assembly [3]. Although these processes are often studied independently, they occur in a highly coordinated manner during protein synthesis [4].

Targeting to the ER is generally mediated by an N-terminal cleavable signal peptide or by the first transmembrane domain of the nascent chain, which can function as a non-cleavable signal-anchor sequence [5]. Importantly, the choice of ER targeting mechanism is closely linked to the topology of the first transmembrane domain [6], [7]. Proteins containing an N-terminal luminal region longer than ∼50 amino acids are more likely to be targeted through a cleavable signal peptide, allowing translocation into the ER lumen prior to membrane insertion. In contrast, proteins with a short or cytosolic N-terminus generally rely on the first transmembrane domain itself as a non-cleavable signal-anchor sequence. This principle underlies the fundamental distinction between signal peptide–driven and signal-anchor–driven insertion pathways.

Interestingly, inspection of multipass ion channels and transporters of the endomembrane system listed in a recent review [8] revealed that the majority do not possess a known or confidently predicted cleavable N-terminal signal peptide, suggesting that they are instead inserted through internal signal-anchor sequences.

TMEM165 is a Golgi-resident membrane protein initially identified in patients affected by congenital disorders of glycosylation (CDG), in which mutations in TMEM165 were reported [9], [10]. It localizes predominantly to the cis-Golgi and exhibits a conserved architecture composed of two inverted repeats of three transmembrane helices connected by a cytosolic loop [11]. Conserved motifs within the first transmembrane helix of each repeat suggest a role in transport activity. TMEM165 has been described as a non-active cation transporter involved in divalent cation homeostasis. Functional studies performed in our laboratory using heterologous yeast and bacterial systems showed that N-terminally truncated variants of human TMEM165 display enhanced functional activity compared with the full-length protein, efficiently complementing the deletion of the yeast orthologue Gdt1 and mediating Ca²⁺ and Mn²⁺ transport in Lactococcus lactis [12]. Subsequent characterization of Gdt1 revealed that Ca²⁺ and Mn²⁺ transport is coupled to H⁺ exchange, leading to the proposal of a reversible Ca²⁺/Mn²⁺– H⁺ antiport mechanism that provides a mechanistic framework for the transport activity of the TMEM165/Gdt1 family [13]. In human cells, calcium and proton transport have been directly demonstrated using electrophysiological approaches [14], [15], whereas a role in manganese homeostasis has been inferred from the restoration of glycosylation defects upon manganese supplementation [16].

Despite recent advances in protein structure prediction, the N-terminal region of TMEM165 remains poorly resolved, with low-confidence structural models [11]. Consequently, two alternative hypotheses have been proposed: the presence of an additional transmembrane segment, or an extended N-terminal region containing a cleavable signal peptide [17], [18]. Resolving this ambiguity is essential for understanding TMEM165 topology and biogenesis. In this study, we sought to determine whether TMEM165 contains a functional signal peptide and to clarify its N-terminal topology.

## Results

### The N-terminal region of TMEM165 is consistent with the presence of a cleavable signal peptide

Structural predictions obtained using AlphaFold3 revealed low confidence for the N-terminal region of TMEM165, preventing a clear conclusion regarding its topology (Figure 1A). Based on previous studies, two models have been proposed: either the presence of an additional transmembrane domain or a long N-terminal tail containing a signal peptide (Figure 1B) [17], [18]. To further investigate this region, we used DeepTMHMM, which predicts both transmembrane domains and protein topology. This analysis identified a putative signal peptide of 33 amino acids with a high confidence score (Figure 1C). Notably, the confidence score associated with this signal peptide prediction is comparable to that obtained for well-characterized proteins such as BiP (Figure 1C) [19], [20]. A conserved motif at positions −3 and −1 relative to the signal peptide cleavage site has previously been reported in large datasets of secreted proteins [21], [22]. This motif is also observed in both BiP and TMEM165 (Figure 1D). To confirm the subcellular localization of endogenous TMEM165, immunofluorescence analysis was performed and showed a clear colocalization with the cis-Golgi marker GM130 (Figure 1E), consistent with previous reports describing TMEM165 as a Golgi resident protein [23].

**Figure 1:**
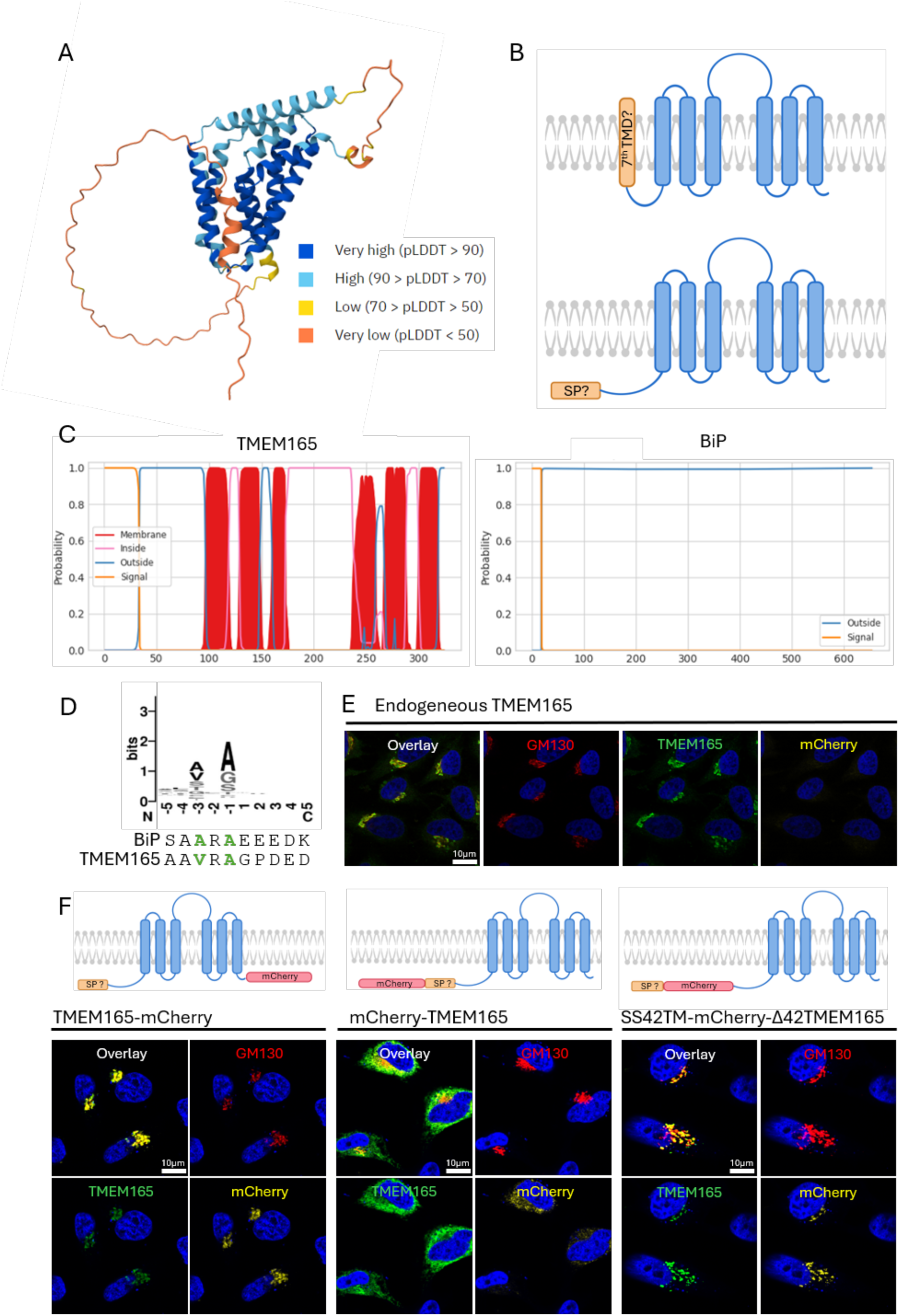
TMEM165 is predicted to contain a cleavable signal peptide. **(A)** AlphaFold was used to predict protein structure; colors correspond to the confidence in the structural prediction of the residues: Orange = very low confidence, Yellow = low, light blue = high and dark blue = very high confidence. In this model, the 84 first amino acids are predicted with very low confidence. **(B)** Two models are described in the literature: the first predicts seven transmembrane domains, while the second includes a cleavable signal peptide. **(C)** Using the DeepTMHMM tool, structural predictions were generated for TMEM165 and for BiP, a known chaperone localized in the ER lumen, by evaluating the probability of each residue belonging to typical protein regions. The yellow line represents the predicted signal peptide, the blue line indicates the extracellular/luminal part of the protein, the pink line corresponds to the intracellular/cytosolic region, and the red columns denote transmembrane domains. **(D)** Top: 1877 secreted protein are aligned to form the sequence logo, comprising 5 amino acids before and after the cleavage site [21]. Bottom: Sequences of BiP chaperone and TMEM165 are compared to the logo. The amino acids forming the cleavage motif are highlighted in green and correspond to the preferred residues at these positions. **(E)** HeLa WT cells fixed and analyzed by fluorescence microscopy. **(F)** HeLa TMEM165 knockout cells were transfected with constructs encoding TMEM165 fused to mCherry at either the N- or C-terminus, or with an additional predicted signal peptide upstream of mCherry. Cells were fixed and analyzed by fluorescence microscopy. DAPI (blue) highlights the nucleus, GM130 (red) marks the Golgi apparatus, TMEM165 is shown in green, and mCherry fluorescence is shown in yellow. C-terminally tagged TMEM165 co-localizes with GM130 (left), whereas N-terminal tagging shows a different distribution (middle). Addition of the predicted signal peptide upstream of mCherry is associated with Golgi co-localization (right).

To experimentally test the functionality of this putative signal peptide, we generated fusion constructs of TMEM165 with the fluorescent protein mCherry. C-terminal tagging did not affect protein localization, as shown by its colocalization with the cis-Golgi marker GM130 (Figure 1F, left panels). In contrast, N-terminal tagging resulted in a clear delocalization of the protein (Figure 1F, middle panels). To determine whether this effect was due to interference with a signal peptide, we engineered a construct in which the predicted signal peptide of TMEM165 was positioned upstream of mCherry, followed by the remainder of the protein. Importantly, nine endogenous amino acids downstream of the predicted cleavage site were included to ensure proper recognition by signal peptidase [21], [24]. This reorganization restored Golgi localization, as indicated by colocalization with GM130 (Figure 1F, right panels). Together, these results support the presence of a functional signal peptide in TMEM165.

### The N-terminal sequence of TMEM165 functions as a cleavable signal peptide

To validate the presence of a functional signal peptide in TMEM165, we assessed its ability to direct a reporter protein into the secretory pathway using both biochemical and imaging approaches. We generated fusion constructs consisting of mCherry alone (mCh), mCherry fused to the nine endogenous amino acids immediately downstream of the predicted TMEM165 cleavage site (9aaTM-mCh), and mCherry fused to the predicted TMEM165 signal sequence (SS) followed by these nine amino acids (SS42TM-mCh) (Figure 2A). Expression of these constructs in HeLa cells was analyzed by immunodetection (Figure 2B). A mobility shift was observed between mCherry alone (mCh) and 9aaTM-mCh, indicating that the addition of these residues could be detected by SDS–PAGE. Importantly, SS42TM-mCh migrated identically to 9aaTM-mCh, consistent with cleavage of the predicted 33-amino-acid signal peptide.

**Figure 2.**
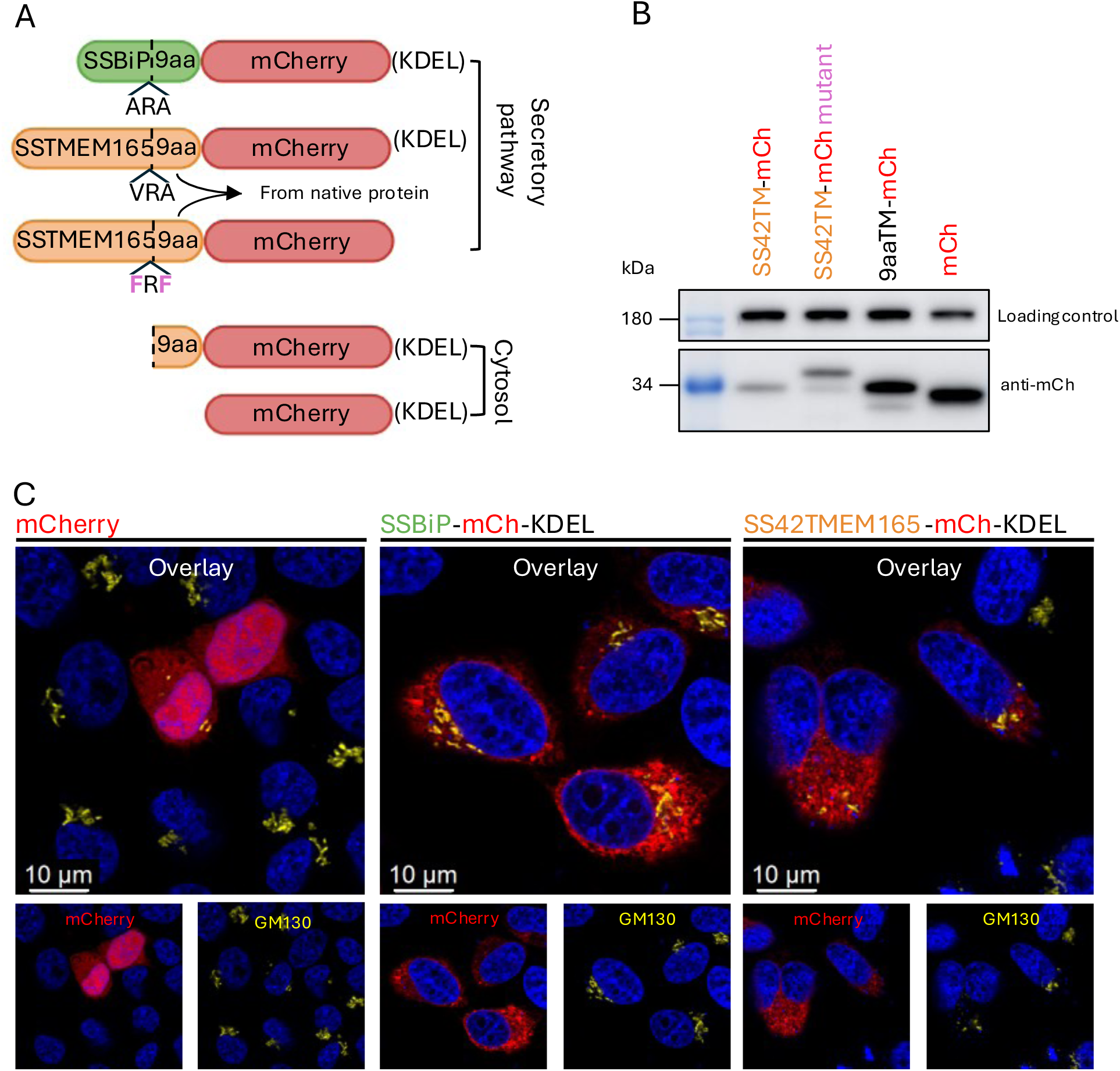
TMEM165 contains a functional cleavable signal peptide that directs mCherry to the secretory pathway. **(A)** Schematic representation of the constructs used: SSBiP–mCherry–KDEL, SS42TM– mCherry–KDEL (with or without VRA→FRF mutation at positions −1 and −3 relative to the predicted cleavage site), 9aaTMEM165–mCherry, and mCherry alone. Constructs containing a signal peptide are expected to enter the secretory pathway, whereas mCherry alone remains cytosolic. The KDEL motif (in parentheses) was included in microscopy constructs to retain proteins in the ER but omitted in samples prepared for size analysis by immunodetection. **(B)** HeLa TMEM165KO cells were transfected with the indicated constructs and lysed 24 h post-transfection. Proteins were separated by SDS-PAGE (13% acrylamide) and analyzed by immunodetection. Clathrin heavy chain was used as a loading control (top), and mCherry was detected using an HRP-conjugated secondary antibody (bottom). SS42TM–mCherry and 9aaTMEM165–mCherry display similar apparent molecular weights, whereas the VRA→FRF mutation in SS42TM–mCherry is associated with the appearance of a higher molecular weight band. **(C)** HeLa cells expressing mCherry (left), SSBiP–mCherry–KDEL (middle), or SS42TM–mCherry–KDEL (right) were fixed 24 h post-transfection and analyzed by confocal microscopy. Nuclei are stained with DAPI (blue), mCherry fluorescence is shown in red, and the Golgi apparatus is labeled with GM130 (yellow). mCherry alone displays a diffuse cytosolic distribution, whereas constructs containing a signal peptide show a reticular and perinuclear pattern.

To disrupt signal peptide cleavage, the residues at positions −3 and −1 were replaced by phenylalanines (VRA→FRF) in the SS42TM-mCh construct (Figure 2A), introducing bulky hydrophobic side chains that are poorly tolerated by signal peptidase [25]. This mutation resulted in the appearance of a higher-molecular-weight form compared to the non-mutated construct, indicating impaired signal peptide cleavage (Figure 2B). However, a residual processed form remained detectable, suggesting incomplete inhibition of cleavage. We next assessed the subcellular localization of these constructs by immunofluorescence. To facilitate the visualization of proteins that had entered the ER, a C-terminal KDEL ER-retention signal was added to all constructs [26]. In the absence of a signal peptide, mCherry exhibited a diffuse cytosolic distribution. In contrast, fusion of the TMEM165 signal sequence yielded a reticular staining pattern, rather than the concentrated perinuclear staining characteristic of the cis-Golgi marker, consistent with localization to the ER. As a positive control, a construct containing the well-characterized BiP signal peptide displayed a similar distribution. This ER-like distribution surrounds the nucleus, consistent with entry into the secretory pathway (Figure 2C).

Similar results were obtained in HEK293T cells (Figure S1A-C, G), in which mCherry alone displayed a diffuse cytosolic distribution, whereas constructs containing either the BiP or TMEM165 signal peptide showed a non-homogeneous, reticular and perinuclear pattern. Together, these results demonstrate that the N-terminal sequence of TMEM165 functions as a cleavable signal peptide capable of directing proteins into the secretory pathway, and that this activity depends on the conserved motif upstream of the cleavage site.

### N-terminal processing determines TMEM165 subcellular localization

To investigate the role of the N-terminal region in TMEM165 targeting, we generated a full-length TMEM165–mCherry construct and two truncation mutants lacking either the signal peptide (Δ33TMEM165–mCherry) or the majority of the N-terminal cytosolic extension (Δ78TMEM165– mCherry) (Figure 1F and 3A). The Δ78 truncation was selected based on a previous study from our laboratory, in which the corresponding construct displayed enhanced functional activity in heterologous systems, efficiently complementing the deletion of Gdt1 in yeast and mediating Ca²⁺ and Mn²⁺ transport in bacterial cells [12]. This construct was also shown to be equivalent to the yeast Δ23Gdt1p variant lacking the putative signal sequence. Subcellular localization was first analyzed in HeLa cells by immunofluorescence. Reference localization patterns corresponding to the Golgi apparatus (GM130), ER (SSBiP-mCh-KDEL), secretory pathway (SSBiP-mCh) and cytosol (mCh) were first established (Figure S2). Based on these reference patterns, transfected cells were classified according to their predominant localization profile (Golgi, ER, secretory pathway, mixed ER/Golgi, or cytosolic). As shown in Figure 3B, full-length TMEM165–mCherry and Δ78TMEM165–mCherry localized predominantly to the Golgi apparatus, whereas Δ33TMEM165–mCherry was mainly detected in the ER. Comparable localization patterns were observed in HEK293T cells (Figure S1D–G).

**Figure 3.**
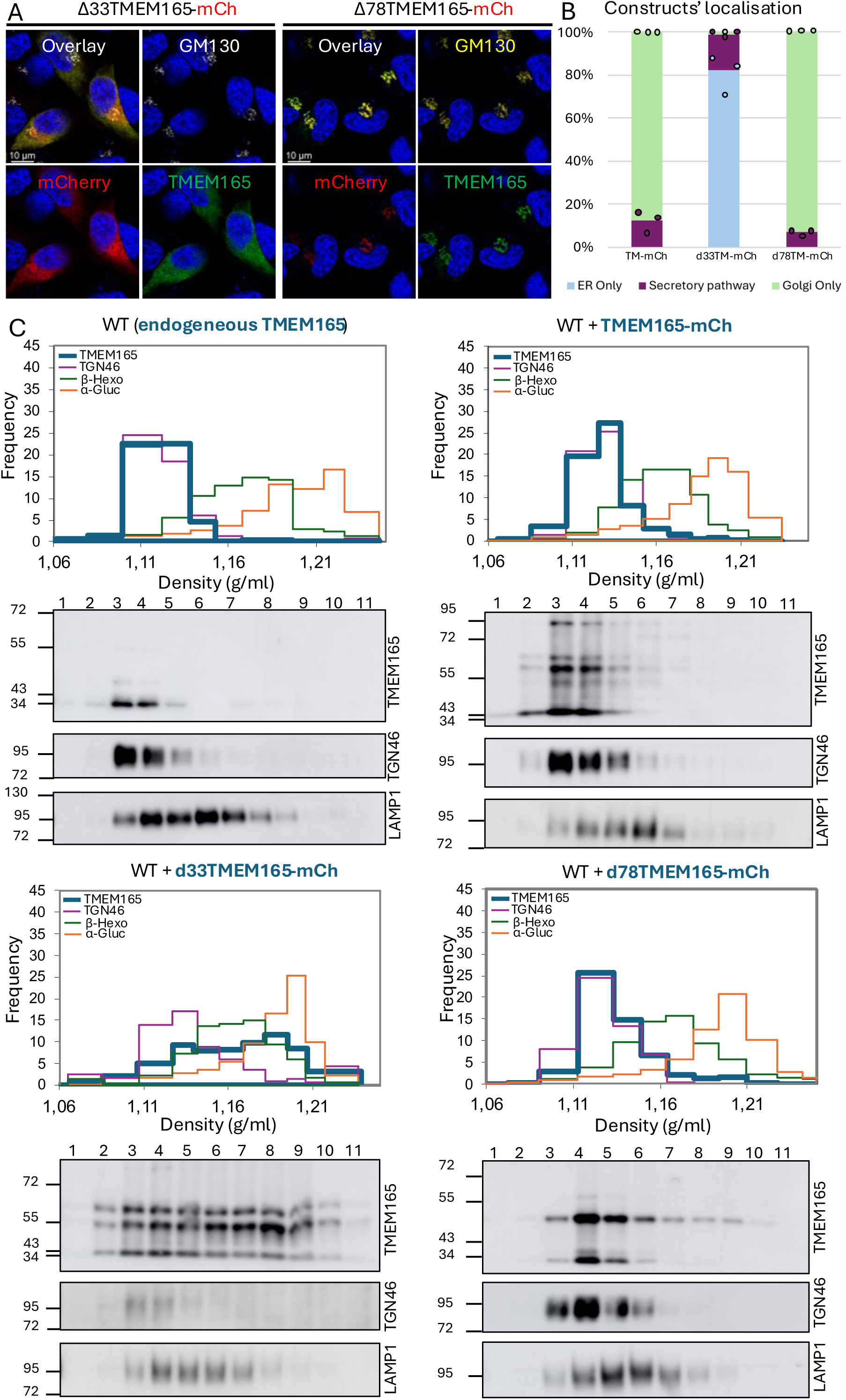
The N-terminal region of TMEM165 controls its subcellular targeting. **(A)** Subcellular localization of TMEM165 variants analyzed by immunofluorescence in HeLa TMEM165KO cells 24 h post-transfection. Cells express Δ33TMEM165–mCherry (left) or Δ78TMEM165–mCherry (right). TMEM165 is shown in green, mCherry fluorescence in red, the Golgi apparatus is labeled with GM130 (yellow/white), and the nucleus are highlighted with DAPI (blue). **(B)** Quantification of subcellular localization patterns from three independent biological replicates. The graph shows the percentage of cells displaying Golgi-only (green), secretory pathway (purple), or ER-only (blue) localization for each construct. Bars represent the mean, and each dot corresponds to one replicate (≥50 cells analyzed per replicate). **(C)** Distribution profiles of endogenous and recombinant TMEM165 variants expressed in HeLa cells after centrifugation of a L and P pooled fraction in a linear sucrose density gradient (see fractionation schematic in Figure S3). Sucrose density (g/mL) is plotted on the x-axis, and the distribution of TMEM165 or organelle markers (normalized to the density increment in each fraction) is plotted on the y-axis. TMEM165 (blue) and TGN46 (purple) distributions were obtained by immunodetection quantification. A lysosomal (β-hexosaminidase, shown in green) and an ER marker (alkaline α-glucosidase, shown in orange) were assessed by enzymatic activity assays. The distribution of a lysosomal transmembrane protein (LAMP1) was also assessed by immunoblotting. For clarity, only one of the lysosomal markers is shown on the graph. (Top left) Endogenous TMEM165. (Top right) Transfected TMEM165–mCherry. (Bottom left) Transfected Δ33TMEM165–mCherry. (Bottom right) Transfected Δ78TMEM165–mCherry. One representative experiment of n=3 independent experiments is shown.

To further validate these observations, subcellular fractionation by differential centrifugation followed by sucrose density gradient separation was performed (Figures S3). Fractions were analyzed by immunodetection and enzyme activity assays using organelle-specific markers for the ER, Golgi apparatus, and lysosomes. Full-length TMEM165–mCherry and Δ78TMEM165–mCherry co-fractionated with the Golgi apparatus marker TGN46, similarly to endogenous TMEM165 (Figure 3C). In contrast, Δ33TMEM165–mCherry exhibited a marked shift in its fractionation profile, largely losing co-fractionation with TGN46 and instead re-distributing towards fractions enriched in ER (α-glucosidase), consistent with the reticular staining pattern observed by fluorescence microscopy and the enrichment of this construct in the microsomal fraction (Figure S4).

Together, these results indicate that the N-terminal signal peptide is required for correct targeting of full-length TMEM165 to the Golgi apparatus, whereas removal of the extended N-terminal region restores Golgi localization. In addition to their distinct localization profiles, full-length and truncated TMEM165 constructs exhibited characteristic migration patterns on immunodetections. Because these patterns suggested the existence of multiple TMEM165 molecular species, they were investigated in greater detail in the next section.

### N-terminal residues of TMEM165 are associated with additional post-translational modifications

The distinct migration patterns observed for full-length and truncated TMEM165 constructs were further investigated by immunodetection analysis. Full-length TMEM165-mCherry was detected as three major species (bands 1–3), whereas Δ33TMEM165-mCherry displayed only bands 2 and 3, and Δ78TMEM165-mCherry migrated as two lower-molecular-weight species (bands 4–5) (Figure 4A–B). These observations suggest that the N-terminal region of TMEM165 is associated with additional post-translational modification events. Sequence analysis of the N-terminal region identified a serine and two threonine residues (S40, T73, T80) located between the signal peptide cleavage site and the first transmembrane domain (Figure 4C), suggesting potential post-translational modifications at these positions. To test this hypothesis, site-directed mutants were generated: S40A, T73A/T80A, and the triple mutant S40A/T73A/T80A in the context of full-length TMEM165-mCherry. Mutation of T73 and T80 resulted in the disappearance of band 1, indicating that this band likely corresponds to a modification dependent on one or both threonine residues (Figure 4B). In contrast, mutation of S40 alone had no detectable effect.

**Figure 4.**
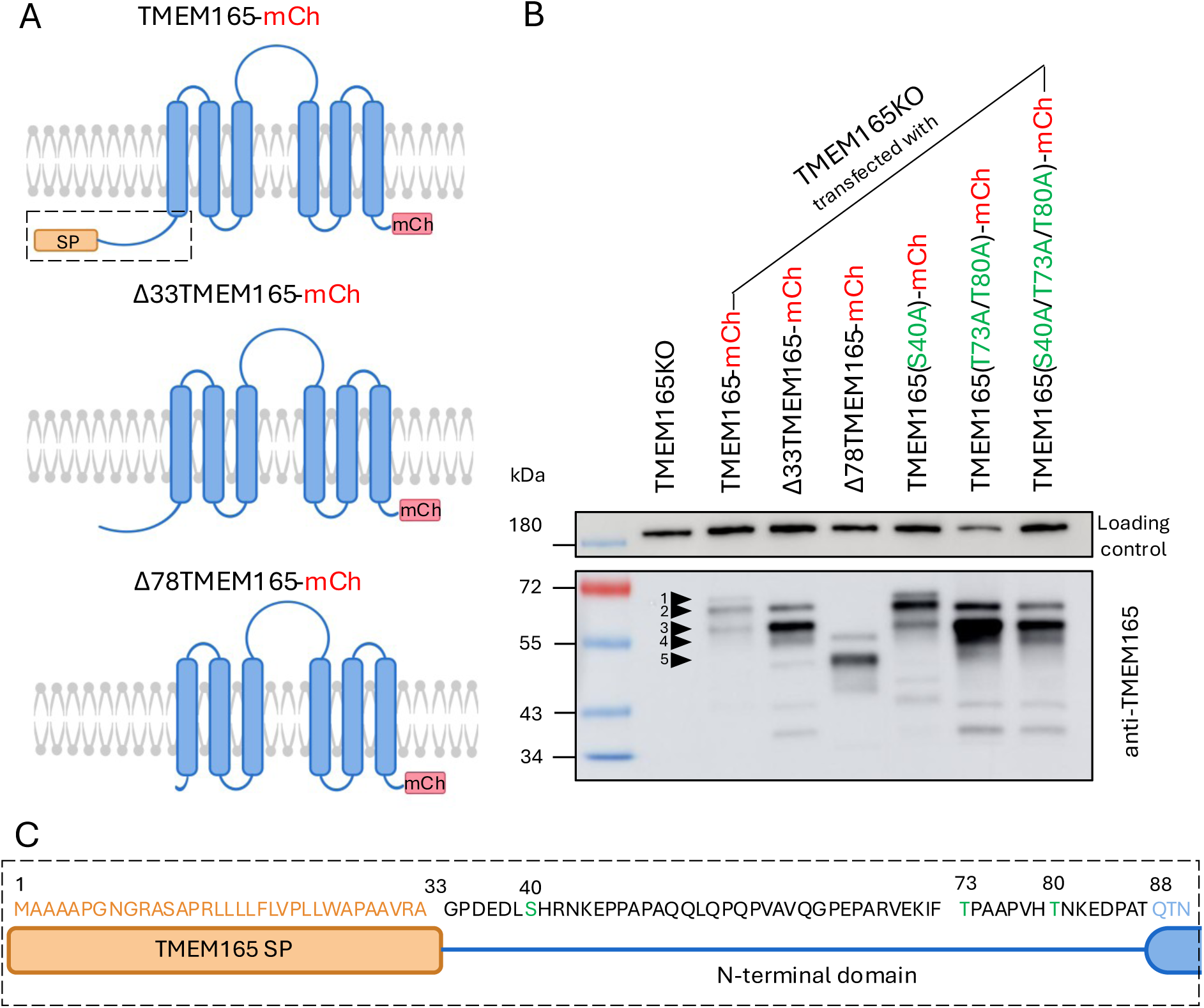
The N-terminal region of TMEM165 undergoes post-translational processing. **(A)** Schematic 2D representations of TMEM165–mCherry constructs illustrating N-terminal truncations: full-length TMEM165–mCherry, Δ33TMEM165–mCherry lacking the signal peptide, and Δ78TMEM165–mCherry lacking the extended N-terminal region. These representations are theoretical and intended to visualize the truncations. **(B)** Immunodetection analysis of TMEM165 variants expressed in TMEM165 knockout HeLa cells. Cells were either non-transfected or transfected with TMEM165–mCherry, Δ33TMEM165–mCherry, Δ78TMEM165–mCherry, or TMEM165–mCherry mutants (S40A, T73A/T80A, S40A/T73A/T80A). Clathrin heavy chain was used as a loading control (top), and TMEM165 was detected using an anti-TMEM165 antibody (bottom). Five distinct bands are indicated by numbered arrows. **(C)** Sequence representation of the N-terminal region of TMEM165 (first 90 amino acids). The predicted signal peptide is highlighted in yellow, and residues targeted for mutagenesis (S40, T73, T80) are shown in green.

Constructs lacking the signal peptide (Δ33TMEM165-mCherry), the extended N-terminal region (Δ78TMEM165-mCherry), or carrying the T73A/T80A mutations all displayed an altered distribution of TMEM165 forms compared with full-length TMEM165-mCherry. Whereas band 2 was the predominant form in the full-length protein, band 3 became the major species in both Δ33TMEM165-mCherry and the T73A/T80A mutant. Similarly, Δ78TMEM165-mCherry exhibited a distinct migration pattern in which band 5 was the predominant form (Figure 4B). These observations suggest that the absence of N-terminal modifications may influence the stability or processing of TMEM165.

### N-terminal truncation alters TMEM165 membrane topology

The observation that deletion of the first 33 amino acids of TMEM165 resulted in ER retention, whereas removal of a larger N-terminal region restored Golgi localization, suggested that these truncations might differentially affect TMEM165 membrane topology. To investigate this possibility, we first performed topology predictions using DeepTMHMM. Predictions were generated for the full-length protein and truncation constructs carrying the HA and mCherry tags introduced to monitor the accessibility of the N- and C-terminal regions in permeabilization assays (Figure S5A-C). Inclusion of these tags did not affect the predicted topology, which remained identical to that of the corresponding untagged sequences. For the Δ33TMEM165 construct, the model predicts a disruption of the first transmembrane segment, resulting in cytosolic exposure of the N-terminus while preserving the overall topology of downstream transmembrane domains (Figure S5A). In contrast, the Δ78TMEM165 construct is predicted to retain a topology similar to full-length TMEM165, with a luminal N-terminus and a cytosolic loop (Figure S5B–C).

To experimentally validate these predictions, we performed epitope accessibility assays using differential membrane permeabilization combined with antibodies directed against the N-terminal HA tag, a cytosolic loop of TMEM165, and the C-terminal mCherry tag. The principle of the assay is summarized in Figure S5D. Selective permeabilization was achieved using streptolysin O (SLO) or digitonin, which permeabilize the plasma membrane while preserving intracellular membranes, whereas Triton X-100 was used to permeabilize all cellular membranes. In Figure 5, only SLO-based experiments are shown for representative imaging, while corresponding digitonin and Triton X-100 datasets are provided in Figure S6 for comparison and validation of permeability conditions. In selective permeabilization experiments, the efficiency of plasma membrane permeabilization was monitored by GM130 immunostaining, as this cytoplasmically exposed Golgi protein is accessible to antibodies only when the plasma membrane has been successfully permeabilized. Quantification was performed on two independent replicates, with at least 50 cells analyzed per replicate.

**Figure 5.**
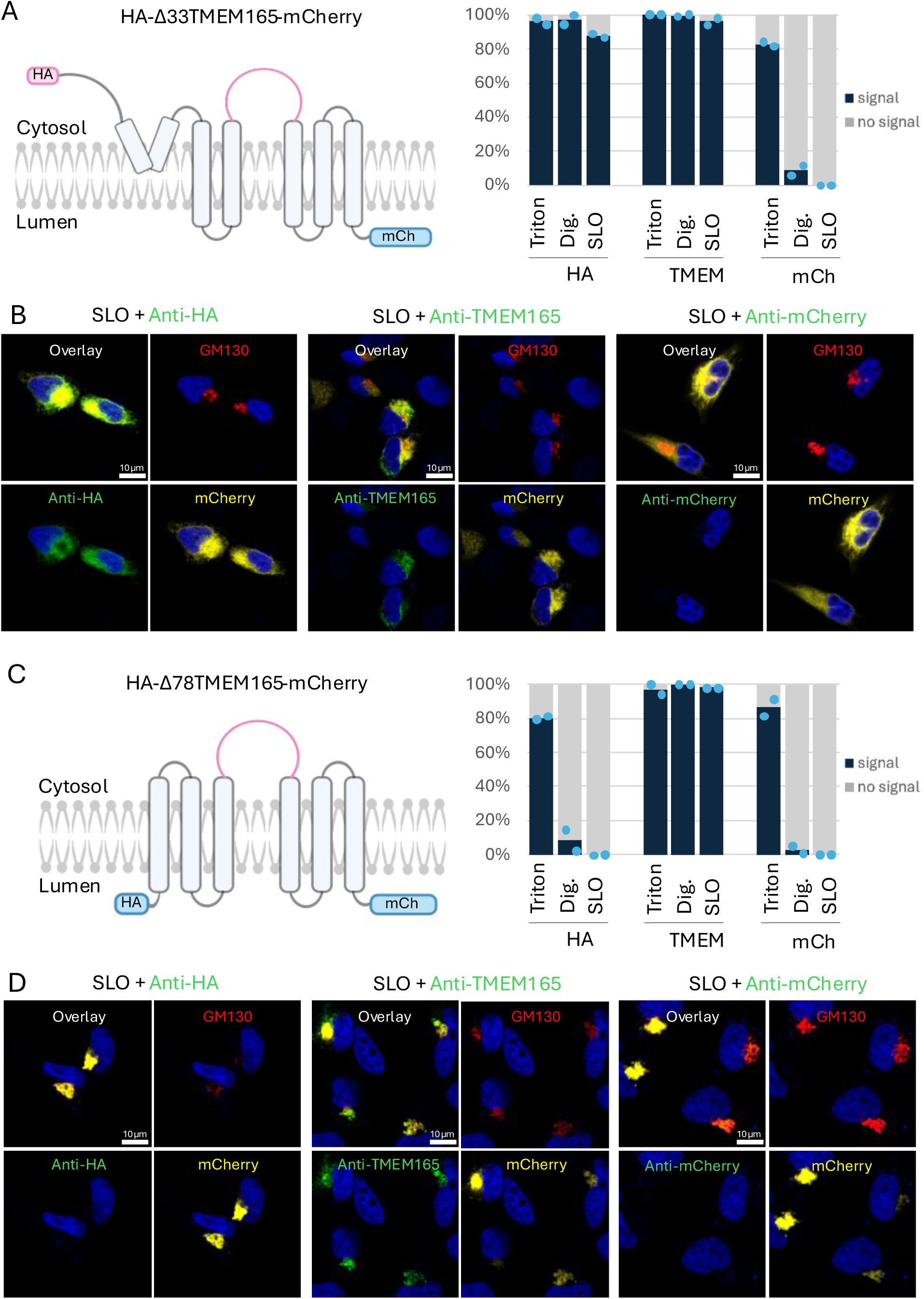
Loss of the signal peptide redirects the TMEM165 N-terminus to the cytosol, while deletion of the extended N-terminal region preserves membrane topology. Schematic representation and membrane permeabilization analysis of HA-Δ33TMEM165-mCherry **(A-B)** and HA-Δ78TMEM165-mCherry **(C-D)** using Triton X-100, digitonin, and streptolysin O (SLO). **(A and C, left)** Two-dimensional model of the predicted topology within the Golgi membrane. Three epitopes were analyzed using specific antibodies: anti-HA recognizing the N-terminal HA tag, anti-TMEM165 recognizing a cytosolic loop of TMEM165, and anti-mCherry recognizing the C-terminal mCherry tag. Pink regions indicate epitopes exposed to the cytosolic side of the membrane, whereas blue regions indicate epitopes located within the Golgi lumen. **(A and C, right)** Percentage of cells displaying antibody staining (“signal”, dark blue) or no detectable staining (“no signal”, gray) following permeabilization with Triton X-100, digitonin, or SLO. Bars indicate the mean percentage obtained from two independent replicates, with a minimum of 50 cells analyzed per replicate. **(B and D)** Representative fluorescence microscopy images obtained after SLO permeabilization. In all conditions, GM130 staining (red) was used as a control for successful plasma membrane permeabilization, while the intrinsic mCherry fluorescence (yellow) allowed identification of transfected cells. Only cells positive for both GM130 and mCherry signals were included in the analysis. The presence or absence of the green signal (corresponding to anti-HA, anti-TMEM165 or anti-mCherry) was used to classify cells into the “signal” or “no signal” categories. Nucleus are highlighted with DAPI (blue). **Left** shows staining with the anti-HA antibody recognizing the N-terminal HA tag. **Middle** shows staining with the anti-TMEM165 antibody recognizing the cytosolic TMEM165 loop. **Right** shows staining with the anti-mCherry antibody recognizing the C-terminal mCherry tag.

Consistent with the predictions, the Δ33TMEM165-mCherry construct showed accessibility of the N-terminal HA epitope under both SLO/digitonin and Triton X-100 conditions, indicating cytosolic exposure (Figure 5A-B). The cytosolic loop was also detected under selective permeabilization conditions, whereas immunodetection of the C-terminal mCherry tag was only observed following Triton X-100 treatment, consistent with a luminal orientation. These results indicate that removal of the first 33 amino acids leads to mislocalization of the N-terminus to the cytosol, while preserving the overall membrane topology of downstream regions.

In contrast, the Δ78TMEM165-mCherry construct exhibited a topology consistent with predictions and full-length TMEM165 organization. Both N-terminal HA and C-terminal mCherry signals were only detectable after Triton X-100 permeabilization, whereas the cytosolic loop was accessible under SLO/digitonin conditions, confirming its cytosolic localization (Figure 5C-D).

Because insertion of an HA epitope at the extreme N-terminus would interfere with signal peptide function, the topology of full-length TMEM165 could not be assessed directly using an N-terminal HA tag. To overcome this limitation, we generated a construct in which the HA epitope was inserted downstream of the signal peptide cleavage site (SS42-HA-d42TMEM165-mCherry), thereby preserving signal peptide-mediated targeting while allowing experimental assessment of N-terminal topology (Figure S7). This construct displayed the expected topology of full-length TMEM165, with both the HA and mCherry epitopes detectable only after Triton X-100 permeabilization, whereas the cytosolic loop remained accessible under selective permeabilization conditions.

Together, these results demonstrate that removal of the signal peptide alters the topology of the N-terminal region of TMEM165. In contrast, the accessibility patterns of the cytosolic loop and C-terminal mCherry tag remained unchanged, indicating that the orientation of downstream domains is preserved. Deletion of the extended N-terminal region preserves the overall membrane orientation of the protein.

### Distinct effects of N-terminal truncations on manganese-induced TMEM165 degradation and glycosylation activity

TMEM165 is a Golgi-resident protein whose stability is tightly regulated by manganese levels and whose activity is required for proper protein glycosylation homeostasis. TMEM165 deficiency has been associated with glycosylation defects affecting highly glycosylated proteins such as LAMP2 [27], whereas manganese exposure triggers a rapid decrease in TMEM165 protein levels through a degradation mechanism that is considered part of its physiological regulation [28]. Together, these readouts provide complementary assays to evaluate TMEM165 regulation and functional activity within the secretory pathway. To further characterize the functional consequences of N-terminal truncations, we first assessed their sensitivity to manganese-induced degradation in HEK293T. Immunodetection analysis revealed that full-length TMEM165-mCherry was efficiently degraded upon manganese treatment, both under transient and stable expression conditions (Figure 6A). A similar degradation pattern was observed for the Δ78TMEM165-mCherry construct. In contrast, the Δ33TMEM165-mCherry construct showed reduced sensitivity to manganese treatment, with no statistically significant decrease in protein levels detected. This behavior was comparable to that of mCherry alone, used as a negative control and not expected to be responsive to manganese.

**Figure 6:**
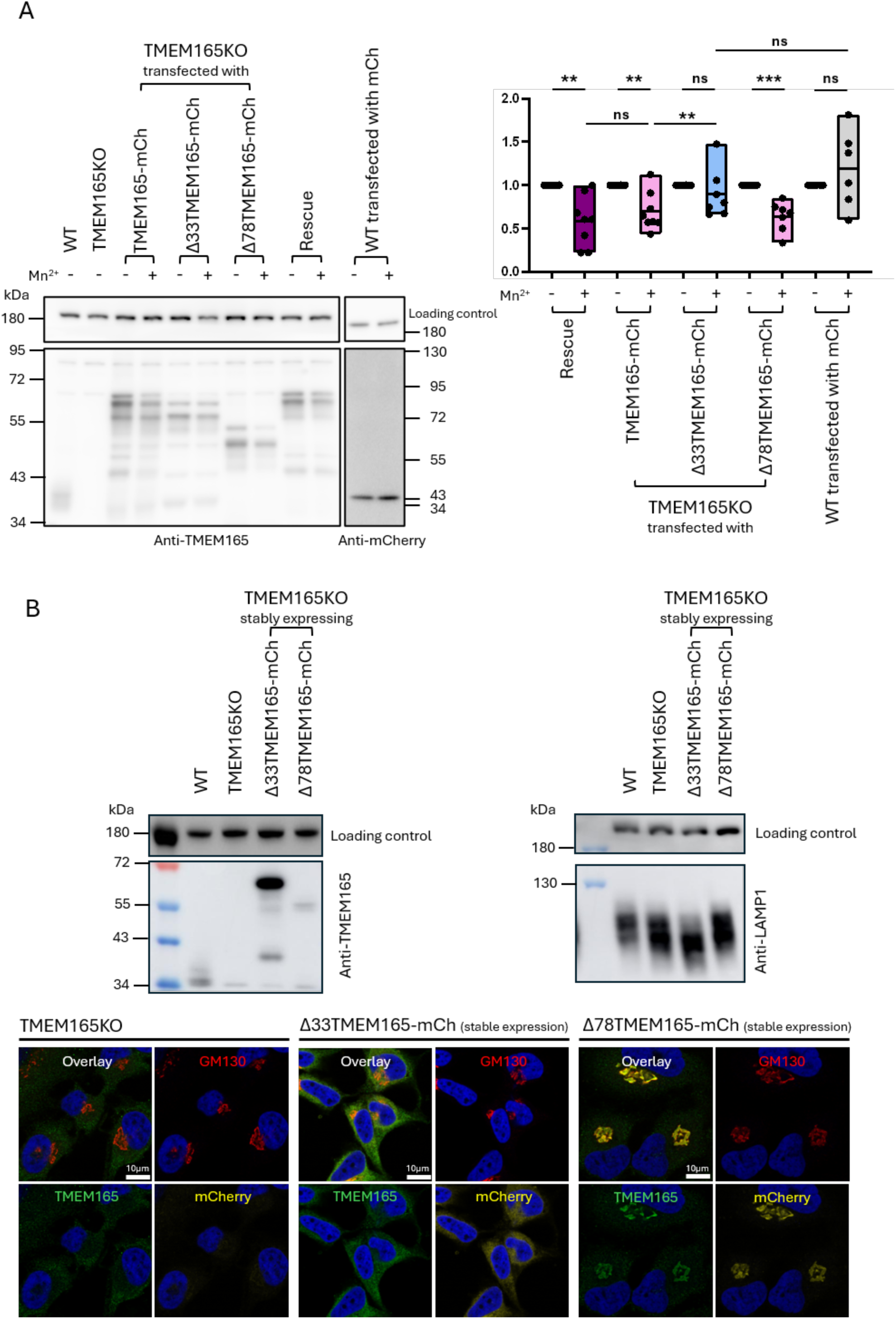
Effect of N-terminal truncations on manganese-induced degradation of TMEM165 and glycosylation activity. **(A)** Immunodetection analysis of HEK293T WT, Rescue, and TMEM165 KO cells transiently expressing full-length TMEM165–mCherry, N-terminal truncation mutants (Δ33TMEM165– mCherry and Δ78TMEM165–mCherry), or mCherry alone (mCh), and treated or not with Mn²⁺. Blots were probed with anti-CHC (loading control), anti-TMEM165, and anti-mCherry antibodies. Quantification of manganese-induced degradation is shown on the right. Signal intensities were normalized to loading control, and the no-Mn²⁺ condition was set as reference (100%), with Mn²⁺-treated samples expressed relative to this baseline [(Mn²⁺ / no Mn²⁺) × 100]. Individual data points represent biological replicates. Statistical significance was assessed using a t-test (ns, not significant; * p < 0.05; ** p < 0.01; *** p < 0.001; **** p < 0.0001). **(B)** Functional analysis of stable HeLa TMEM165 KO cell lines stably expressing Δ33TMEM165–mCherry or Δ78TMEM165–mCherry. **Top:** Immunodetection of TMEM165 and LAMP1 in stable HeLa WT, KO, Δ33TMEM165–mCherry, and Δ78TMEM165–mCherry cell lines, with clathrin heavy chain (CHC) used as loading control. **Bottom:** Representative fluorescence microscopy images of the same stable HeLa cell lines analyzed above (KO, Δ33TMEM165–mCherry, and Δ78TMEM165–mCherry; left to right). Cells were fixed and analyzed by fluorescence microscopy. DAPI (blue) highlights nuclei, GM130 (red) marks the Golgi apparatus, TMEM165 is shown in green, and mCherry fluorescence is shown in yellow.

We next investigated whether these constructs retained TMEM165 functional activity by assessing their ability to rescue the glycosylation defects observed in TMEM165-deficient cells. As previously reported, highly glycosylated proteins such as LAMP2 display increased electrophoretic mobility in TMEM165-deficient cells, reflecting their reduced glycosylation status. To this end, stable HeLa TMEM165KO cell lines expressing either Δ33TMEM165-mCherry or Δ78TMEM165-mCherry were generated. Importantly, both immunodetection and immunofluorescence analyses confirmed that these stable cell lines recapitulated the expression levels and subcellular localization patterns observed under transient conditions, with Δ33TMEM165-mCherry remaining mislocalized and Δ78TMEM165-mCherry maintaining Golgi localization (Figure 6B).

Functional analysis revealed that expression of Δ33TMEM165-mCherry failed to rescue the LAMP2 glycosylation defect and instead resulted in an even more pronounced alteration of its electrophoretic mobility compared with TMEM165KO cells (Figure 6B). In contrast, stable expression of Δ78TMEM165-mCherry restored, at least partly, a normal LAMP2 migration profile, indicating that this truncated form retains functional activity despite the absence of the extended N-terminal region.

Together, these results demonstrate that Δ78TMEM165-mCherry remains responsive to manganese-induced degradation and partially restores glycosylation despite the absence of the extended N-terminal region. In contrast, Δ33TMEM165-mCherry, previously shown to display altered topology and aberrant localization, exhibits reduced sensitivity to manganese-induced degradation and fails to rescue TMEM165-dependent glycosylation defects. These findings indicate that the extended N-terminal region is dispensable for TMEM165 function, whereas proper topology and targeting of the protein are essential for its physiological activity.

## Discussion

Our study reveals that TMEM165, a multi-pass Golgi membrane protein, relies on a functional N-terminal signal peptide for correct targeting and membrane topology. This conclusion is supported by converging bioinformatic and experimental evidence. In its absence, full-length TMEM165 is mislocalized, as shown by immunofluorescence and fractionation analyses revealing loss of Golgi association (TGN46/GM130) and redistribution toward ER- and lysosomal compartments. Topological analyses further show that signal peptide deletion selectively disrupts N-terminal organization, leading to cytosolic exposure of the N-terminus while downstream regions, including the central loop and C-terminus, retain their expected orientation, suggesting that the primary defect affects early membrane insertion events involving the first transmembrane segment, in agreement with predictions. In contrast, deletion of a larger N-terminal region restores Golgi targeting and preserves overall topology, indicating that the first transmembrane helix can act as a signal-anchor sequence, as described for other multi-pass proteins [6], whereas the conservation of a cleavable signal peptide suggests an additional role for the luminal N-terminal region.

A striking feature of TMEM165 and its orthologues is the remarkable evolutionary diversity of the N-terminal region. It is absent from bacterial orthologues, progressively expands in prokaryotes and eukaryotes, and is replaced in plant orthologues (PAM71 and CMT1) by predicted and experimentally supported chloroplast transit peptides [11], [29], [30]. This diversification suggests that N-terminal targeting sequences have evolved in parallel with increasing cellular compartmentalization. Importantly, our data indicate that neither the signal peptide nor the extended N-terminal region is strictly required for the core transport activity of the protein. Previous studies demonstrated that N-terminally truncated versions of human TMEM165 (Δ78), Arabidopsis PAM71 (Δ155) and Arabidopsis CMT1 (Δ131) remain functional when expressed in yeast, where they complement the calcium-sensitive phenotype associated with GDT1 deficiency [12], [31], [32]. In the present study Δ78TMEM165 exhibits proper Golgi localization, membrane topology, manganese-induced degradation, and partially restores the glycosylation defects associated with TMEM165 deficiency. Together, these observations provide evidence that the extended N-terminal region is dispensable for TMEM165 function in human cells and argue against a direct role of this domain in ion transport itself, raising the question of why it has been acquired and conserved during evolution.

The present findings draw particular attention to the region located between the signal peptide cleavage site and the first transmembrane domain of TMEM165 Although dispensable for the core transport activity and manganese-dependent degradation examined here, this segment has been introduced and maintained during evolution, suggesting the existence of an additional functional constraint acting on this region. Notably, it contains a cluster of acidic (Asp36, Glu37, Asp38 and Glu45) and basic (His41, Arg42 and Lys44) residues as well as several serine and threonine residues that could represent sites of post-translational modification [17]. The charged region is reminiscent of regulatory domains described in other Ca²⁺/H⁺ antiporter families. However, unlike the N-terminal autoinhibitory domains of transporters such as CAX1 [33], [34], this region is predicted to reside in the lumen rather than the cytosol, suggesting a distinct mode of regulation. Interestingly, mutation of Thr73 and Thr80 abolishes higher molecular weight species detected by immunodetection (Figure 4). Combined with our topological data, this observation is more consistent with a luminal post-translational modification such as O-glycosylation than with cytosolic phosphorylation. Indeed, as established in standard glycobiology references, O-linked glycosylation of serine and threonine residues in secretory and membrane proteins occurs within the lumen of the ER and Golgi apparatus, whereas serine/threonine phosphorylation is mediated by cytosolic and nuclear kinases [35]. Although the precise function of this region remains unknown, its evolutionary conservation, particular composition, and apparent susceptibility to post-translational modification strongly suggest that it contributes to TMEM165 biogenesis or regulation.

Direct comparisons with other multi-pass transporters remain limited, as inspection of multi-pass ion channels and transporters of the endomembrane system listed in a recent review [8] revealed that the majority do not possess a known or confidently predicted cleavable N-terminal signal peptide. A more informative framework can be found in G protein-coupled receptors (GPCRs), which also possess long luminal or extracellular N-terminal domains and have been extensively studied in terms of N-terminal biogenesis and regulation [36]. A subset of GPCRs contains cleavable signal peptides whose presence cannot be explained solely by topological requirements, suggesting additional roles in protein biogenesis beyond membrane targeting. Consistent with this idea, GPCR N-terminal domains are frequently subject to post-translational modifications that regulate folding, trafficking and stability [37].

In this context, the present data support a model in which the TMEM165 signal peptide ensures efficient translocation of the N-terminal region into the ER lumen, thereby promoting the establishment of the correct membrane topology required for ER exit and Golgi localization. In the absence of signal peptide-mediated translocation, the N-terminus remains exposed to the cytosol, resulting in an altered topology that correlates with ER retention and loss of Golgi targeting.

However, the mechanism underlying this requirement remains unclear. One possibility is that efficient luminal translocation requires signal peptide–mediated entry of the relatively long N-terminal region, as previously suggested for other multi-pass membrane proteins [7]. Alternatively, signal peptide cleavage may be required to ensure correct TMEM165 biogenesis by preventing cytosolic exposure of the N-terminal region, which could otherwise undergo post-translational modifications that interfere with proper topology establishment. In line with this possibility, ribosome-associated kinases have been implicated in co-translational phosphorylation events on emerging nascent chains [38]. According to the positive-inside rule, membrane topology is primarily influenced by the distribution of positively charged residues across the membrane [39], suggesting that additional negative charges introduced by phosphorylation would not be expected to act as primary topological determinants. However, phosphorylation has been shown to dynamically regulate membrane protein topology in a lipid-dependent manner, indicating that its effects may extend beyond simple electrostatic considerations [40]. Nevertheless, whether similar mechanisms contribute to TMEM165 biogenesis remains unknown.

At this stage, we cannot distinguish between these possibilities. However, our findings indicate that the cleavable signal peptide is required for correct N-terminal translocation, membrane topology, and Golgi targeting of TMEM165.

## Materials & Methods

### Antibodies and reagents

The following antibodies were purchased from the indicated suppliers: mouse anti-clathrin heavy chain (BD Transduction Laboratories, 610500, 1:3000 for immunodetection); mouse anti-GM130 (BD Transduction Laboratories, 610822, 1:200 for immunofluorescence); rat anti-HA high affinity (Roche, 11867423001, 1:100 for immunofluorescence); rabbit anti-LAMP1 (Cell Signaling Technology, 9091, 1:1000 for immunodetection); mouse anti-LAMP2 (Santa Cruz Biotechnology, sc-18822, 1:1000 for immunodetection); rabbit anti-mCherry (Rockland, 600-401-P16S, 1:3000 for immunodetection and 1:250 for immunofluorescence); rabbit anti-TGN46 (Invitrogen, PA5-23068, 1:1000 for Immunodetection); rabbit anti-TMEM165 (Proteintech, 20485-1-AP, 1:2000 for immunodetection and 1:800 for immunofluorescence).

Secondary antibodies used were goat anti-mouse IgG HRP-conjugated secondary antibody (Chemicon International, AP308P, 1:10000 for immunodetection chemiluminescence); goat anti-rabbit immunoglobulins/HRP (Dako, P0448, 1:5000 for immunodetection chemiluminescence); Alexa Fluor 647 goat anti-mouse IgG (H+L) (Invitrogen, A21236, 1:250 for immunofluorescence); Alexa Fluor 488 goat anti-rabbit IgG (H+L) (Invitrogen, A11034, 1:250 for immunofluorescence); and Alexa Fluor 488 goat anti-rat IgG (H+L) (Invitrogen, A11006, 1:250 for immunofluorescence).

### Plasmid constructs and mutagenesis

All constructs used in this study were generated in the pIRES-neo expression vector backbone. TMEM165-derived deletion, tagging, signal peptide, retention signal, and point-mutant constructs were generated by a combination of amplification-restriction-ligation cloning, partial plasmid amplification, sequence insertion using primers containing 5′ overhangs, site-directed mutagenesis, and triple PCR approaches. A complete list of constructs, parental plasmids, cloning strategies, and primer sequences is provided in Table S1. All constructs were verified by Sanger sequencing. The pIRES-neo empty vector, pIRES-neo-mCherry, and pRS416-pTPI-TPARL plasmids were used as starting materials. The sequence encoding the first 27 amino acids of BiP was synthesized by GenScript (Piscataway, NJ, USA) and delivered in a pUC57 plasmid. All cloning procedures were performed using chemically competent E. coli TOP10 cells.

### Cell culture and transfection

HeLa wild-type (WT) and TMEM165 knockout (TMEM165KO) cell lines were kindly provided by Marja Jäättelä (Copenhagen, Denmark) [14]. HEK293T WT and TMEM165KO cell lines were kindly provided by François Foulquier (Lille, France) [10]. HeLa and HEK293T cells were maintained at 37°C in a humidified atmosphere containing 5% CO₂ in high-glucose DMEM medium supplemented with 10% fetal bovine serum (PAN Biotech, #P30-3305), 1% penicillin-streptomycin, and 1 mM sodium pyruvate. For transfection of HeLa cells, 6 × 10⁴ cells per well were seeded in 24-well plates 24 h before transfection. The next day, the culture medium was replaced with transfection medium containing 50 µL Opti-MEM (Gibco, #31985062), 0.6 µL FuGENE reagent (Promega, #E2311), and 200 ng plasmid DNA, completed to a final volume of 500 µL with DMEM. For transfection of HEK293T cells, cells were seeded simultaneously with the transfection mixture in 24-well plates. Cell suspension was prepared from a flask at approximately 90% confluence by diluting 5 mL of cell suspension into 20 mL of fresh medium. Then, 750 µL of diluted cells were added per well together with 100 µL of transfection mix containing Opti-MEM, plasmid DNA, and PEI (Tocris, #7854). The amounts of DNA and PEI were determined using the Cytographica PEI Transfection Calculator available online. HeLa and HEK293T cells were harvested 24 h after transfection for subsequent experiments.

### Generation of stable cell lines

Stable HeLa TMEM165KO cell lines expressing Δ33TMEM165-mCherry or Δ78TMEM165-mCherry were generated by nucleofection of the corresponding pIRES-neo expression constructs. Following nucleofection, cells were allowed to recover for several days before the addition of G418 (Geneticin) at a concentration of 280 µg/mL for selection. Polyclonal resistant populations were subsequently expanded and maintained under continuous selection. Expression and subcellular localization of the transgenes were verified by immunodetection and immunofluorescence prior to further analyses.

### Cell lysis

For whole-cell extracts from transfected cells grown in 24-well plates, confluent cells were placed on ice, washed once with PBS, and lysed directly in Laemmli sample buffer containing Tris-HCl pH 6.8, SDS, glycerol, bromophenol blue, and β-mercaptoethanol. Samples obtained from subcellular fractionation and sucrose gradient experiments were mixed with 4× Laemmli sample buffer containing 250 mM Tris-HCl pH 6.8, 8% SDS, 40% glycerol, 0.04% bromophenol blue, and 5% β-mercaptoethanol. In all cases, samples were heated at 95°C for 10 min before analysis.

### Immunodetection analysis

Proteins were separated by SDS-PAGE using home-made polyacrylamide gels composed of a stacking gel and an 8% resolving gel. Electrophoresis was performed in Tris-glycine-SDS running buffer at 100 V for 30 min followed by 140 V until the migration front reached the bottom of the gel. For experiments requiring improved resolution of glycosylation shifts, migration time was extended. Proteins were transferred onto membranes using Transfer Pack membranes (Bio-Rad, #1704156) at 1.3 A with a maximum voltage of 25 V for 10 min. Membranes were blocked in PBS containing 5% (w/v) milk powder and 0.1% Tween-20 and incubated with primary and secondary antibodies diluted in the same buffer. Signal detection was performed by chemiluminescence using SuperSignal West Femto substrate (Thermo Scientific, #34095) and acquisition with an Amersham Imager 600 (GE Healthcare).

### Immunofluorescence microscopy

Cells grown on coverslips were washed with PBS supplemented with MgCl₂ and CaCl₂ (PBS++) and fixed with 4% paraformaldehyde for 15 min at 37 °C. Residual paraformaldehyde was quenched with NH₄Cl before permeabilization in PBS containing 1% BSA and 0.1% saponin. Primary antibodies were diluted in permeabilization buffer and incubated on coverslips for 30 min at room temperature. After washing, coverslips were incubated with Alexa Fluor-conjugated secondary antibodies under the same conditions. Coverslips were mounted using Fluoromount-G containing DAPI. Images were acquired using a Leica Stellaris 8 Falcon confocal microscope at a resolution of 1024 × 1024 pixels. For each condition, at least 50 transfected cells were analyzed.

### Subcellular fractionation

HeLa cells were fractionated into six fractions (E, extract or post-nuclear supernatant; N, nuclear fraction; M, heavy mitochondrial fraction; L, light mitochondrial fraction; P, microsomal fraction; and S, soluble fraction) by differential centrifugation following a slightly modified version of de Duve’s fractionation scheme [41], [42]. The pooled L and P fractions were then centrifuged in a linear sucrose density gradient ranging from 1.09 to 1.34 g/mL for 16h at 35 000 RPM in a SW5Ti rotor. The collected fractions were analyzed by immunodetection or enzymatic assays, as described in the corresponding sections.

### Enzyme activity assays

The activities of alkaline α-glucosidase (ER marker) and β-hexosaminidase (lysosomal marker) were measured as described [42].

### Manganese treatment and degradation assays

HEK293T cells were treated with 100 µM MnCl₂ (final concentration) added at the time of cell seeding [28]. Cells were harvested the following day for analysis. For transient transfections, PEI-mediated transfection was performed concomitantly with MnCl₂ treatment. For all conditions, cells were lysed and proteins were analyzed by immunodetection as described above. Quantification of manganese-induced degradation was performed by densitometric analysis of immunodetection bands. Signal intensities were normalized to a loading control, and the no-Mn²⁺ condition was set as reference (100%). Mn²⁺-treated samples were expressed relative to this baseline using the formula: (Mn²⁺ / no Mn²⁺) × 100.

### Differential permeabilization assays

Selective membrane permeabilization experiments were performed on transfected HeLa cells grown on coverslips and were adapted from previously described digitonin-based topology assays used for TMEM165 characterization [28], with additional permeabilization using streptolysin O (SLO), a cholesterol-dependent pore-forming toxin that selectively permeabilizes the plasma membrane while preserving intracellular membrane integrity [43], [44]. Then, three permeabilization conditions were used: digitonin (30 µg/mL), streptolysin O (SLO; 0.5 U/mL, activated with 10 mM DTT from a 100 U/mL stock), or Triton X-100 (0.1%). Cells were washed with PBS prior to treatment. Digitonin and SLO treatments were performed at 4 °C, whereas Triton X-100 treatment was performed at room temperature. Cells were incubated with digitonin for 15 min or with activated SLO for 10 min, followed by two washes with cold PBS. Cells were then fixed with 4% paraformaldehyde prewarmed at room temperature for 15 min, followed by three washes with PBS. For complete permeabilization controls, cells were incubated with 0.1% Triton X-100 for 10 min at room temperature after fixation and washed three times with PBS. Cells were blocked for 1 h at room temperature in blocking buffer composed of gelatin (0.1% w/v), BSA (2% w/v), and FBS (2% v/v) in PBS. Primary antibodies were incubated for 1 h at room temperature, followed by three washes with PBS. Secondary antibodies were then incubated for 1 h at room temperature in the dark, followed by three additional washes with PBS. Coverslips were mounted and imaged as described for immunofluorescence microscopy.

### Topology prediction

Protein topology was predicted using DeepTMHMM via the online server (https://dtu.biolib.com/DeepTMHMM/). Predictions were performed using default parameters.

## Acknowledgments

The authors would like to thank Martine Albert and Florentine Gilis for their valuable technical assistance. The authors also thank Majda Belataris and Loïc Fangang Fanseu for their contribution to the generation of experimental replicates. The authors gratefully acknowledge Marja Jäättelä and François Foulquier for providing the cell lines used in this study.

## Supplementary Material

**Figure S1.**
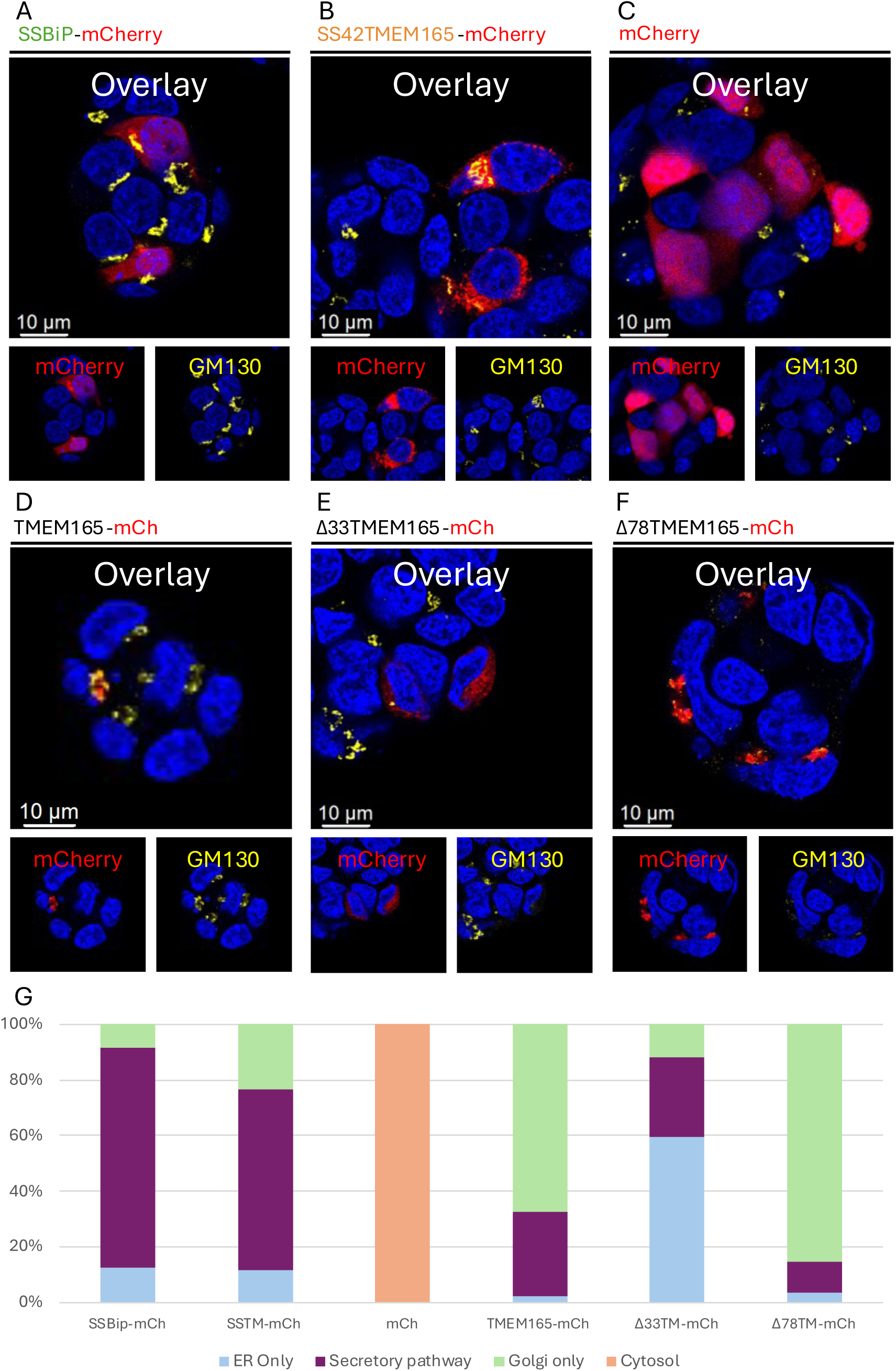
TMEM165 localization patterns are conserved in HEK2G3T cells. **(A–F)** HEK293T cells were transfected with the indicated constructs, fixed 24 h post-transfection, and analyzed by fluorescence microscopy. Cells express SSBiP–mCherry (A), SS42TMEM165–mCherry (B), mCherry alone (C), TMEM165– mCherry (D), Δ33TMEM165–mCherry (E), or Δ78TMEM165–mCherry (F). mCherry fluorescence is shown in red, the Golgi apparatus is labeled with GM130 (yellow), and nucleus are highlighted with DAPI (blue). **(G)** Quantification of subcellular localization patterns based on the analysis of at least 30 cells per condition. Cells were classified into four categories: “ER only” (reticular and perinuclear signal), “Golgi only” (perinuclear punctate signal co-localizing with GM130), “secretory pathway” (combined reticular/perinuclear and Golgi-associated signals), and “cytosol” (diffuse signal throughout the cell).

**Figure S2.**
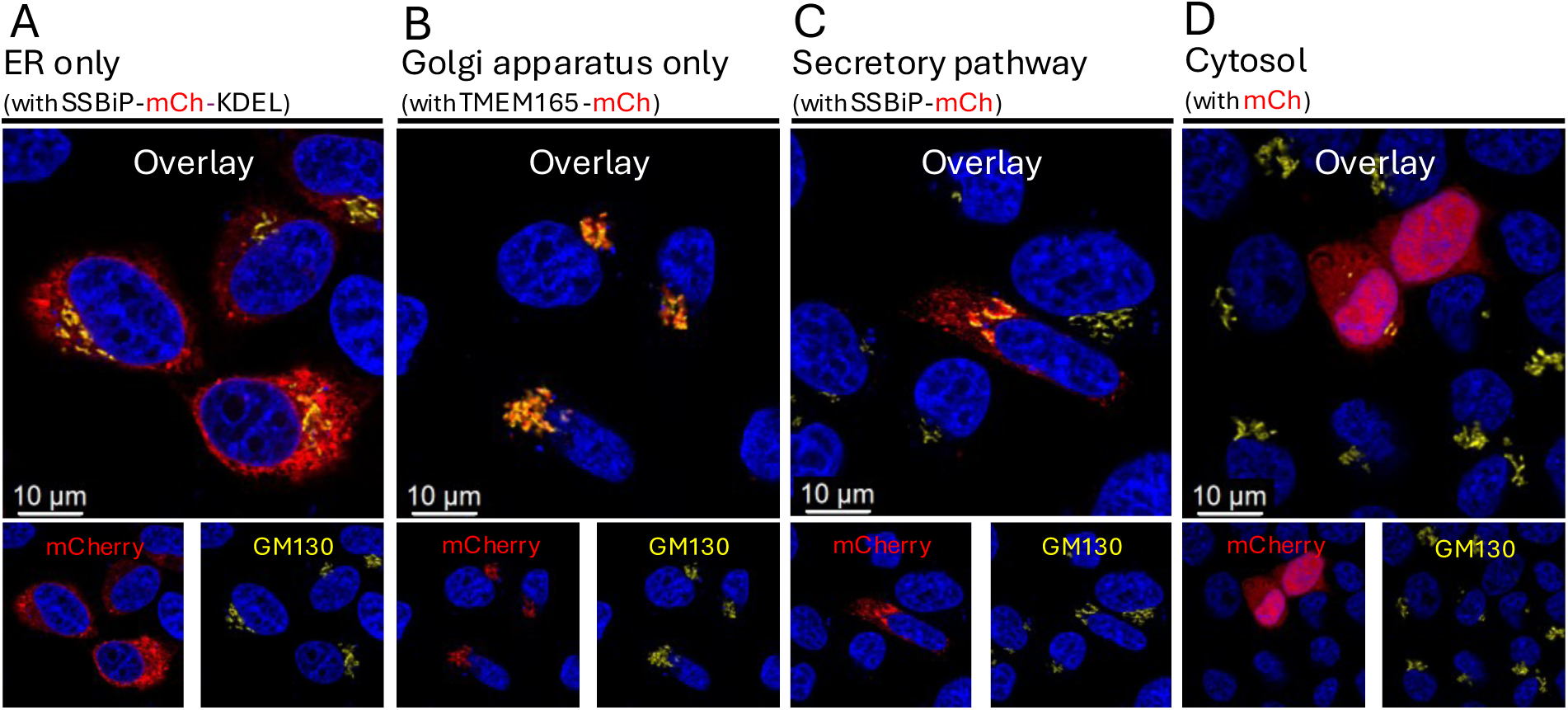
Reference patterns used for subcellular localization classification. HeLa cells were transfected with the indicated constructs, fixed 24 h post-transfection, and analyzed by fluorescence microscopy to define reference localization patterns. mCherry fluorescence is shown in red, the Golgi apparatus is labeled with GM130 (yellow), and nucleus are highlighted with DAPI (blue). **(A)** ER localization is shown using SSBiP–mCherry–KDEL, displaying a reticular and perinuclear distribution. **(B)** Golgi localization is defined by GM130 immunostaining, showing a punctate perinuclear signal. **(C)** Secretory pathway localization is represented by SSBiP–mCherry, showing combined ER-like and Golgi-associated patterns. **(D)** Cytosolic localization is shown using mCherry alone, displaying a diffuse distribution throughout the cell.

**Figure S3.**
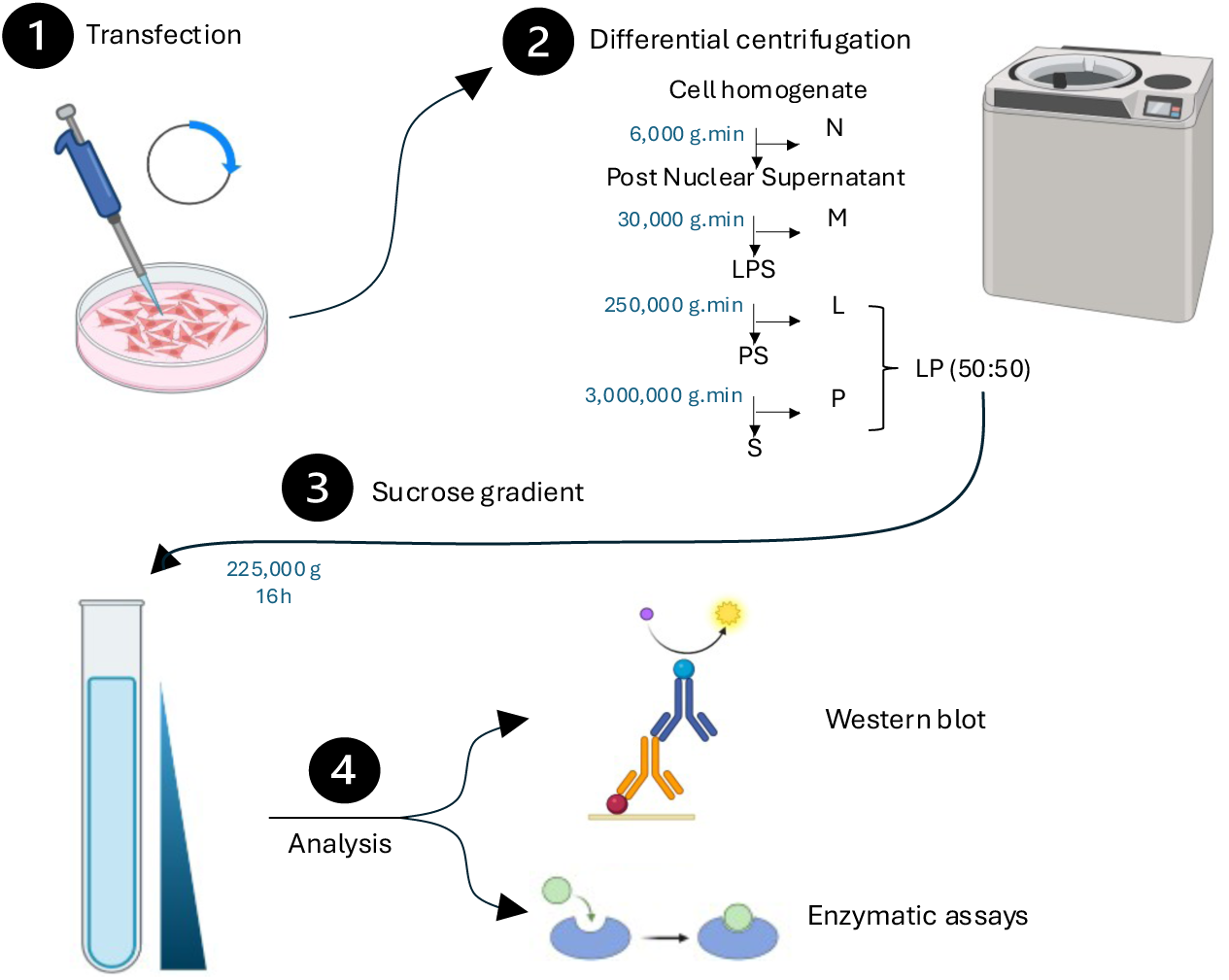
Experimental workflow for subcellular fractionation and sucrose density gradient analysis. HeLa cells were homogenized and subjected to sequential differential centrifugation to obtain the nuclear (N), heavy mitochondrial (M), light mitochondrial (L), microsomal (P), and soluble (S) fractions from the post-nuclear supernatant (E). The pooled L and P fractions were subsequently separated on a linear sucrose density gradient (1.09–1.34 g/mL) by ultracentrifugation. Gradient fractions were collected and analyzed by immunodetection and enzymatic activity assays.

**Figure S4.**
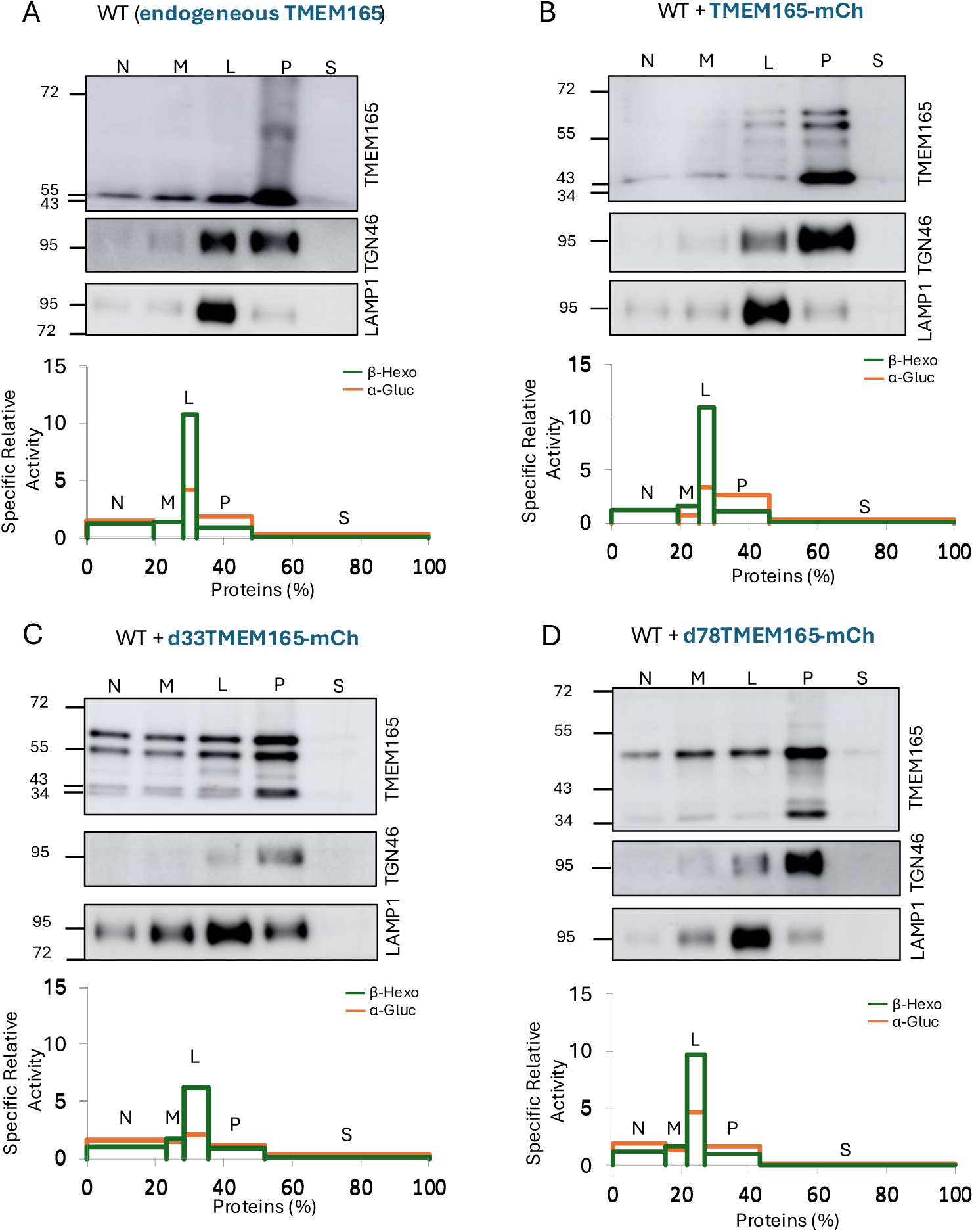
Subcellular fractionation of TMEM165 variants and characterization of organelle-enriched fractions. Immunodetection analysis of subcellular fractions obtained after differential centrifugation of a HeLa cells homogenate (transfected of not with the indicated constructs) as described in Figure S3. Following removal of the nuclear fraction (N), the post-nuclear extract (E) was subjected to sequential centrifugation steps to generate the mitochondrial (M), light mitochondrial (L), microsomal (P; enriched in ER and Golgi membranes), and soluble (S) fractions. Fractions were analyzed by immunodetection using antibodies against TMEM165, TGN46 (Golgi marker), and LAMP1 (lysosomal marker) Equal amounts of proteins (2 µg per fraction) were loaded for each fraction. Graphs shown below each blot represent the specific relative activities of β-hexosaminidase (lysosomal marker, shown in green) and alkaline α-glucosidase (ER marker, shown in orange) as a function of the percentage of total proteins recovered in each fraction. **(A)** Endogenous TMEM165 distribution. **(B)** Distribution of transfected TMEM165–mCherry. **(C)** Distribution of transfected Δ33TMEM165–mCherry. **(D)** Distribution of Δ78TMEM165–mCherry.

**Figure S5.**
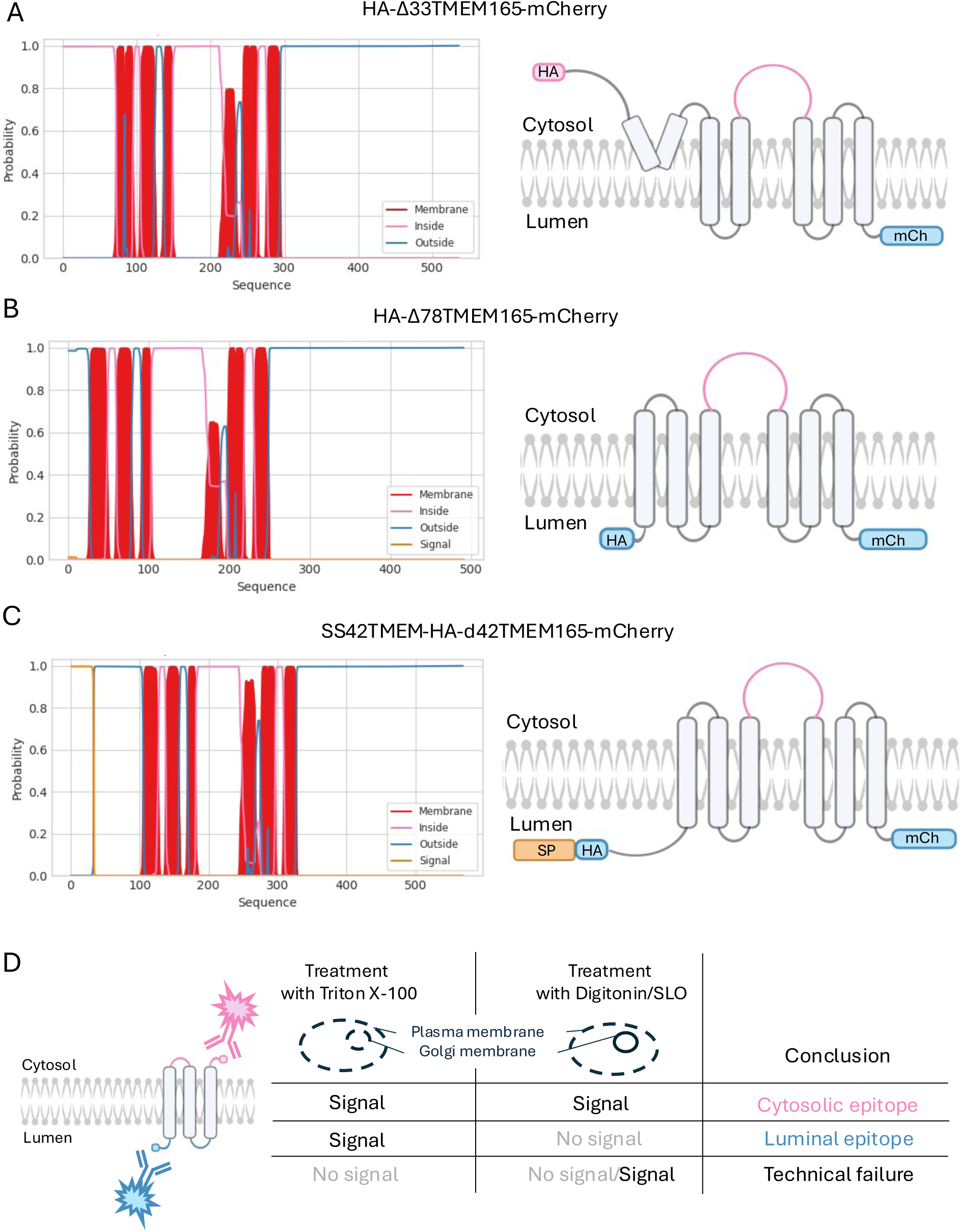
Topology prediction of TMEM165 constructs using DeepTMHMM and schematic overview of the permeabilization-based topology assay. Predicted membrane topology of HA-Δ33TMEM165-mCherry **(A)**, HA-Δ78TMEM165-mCherry **(B)**, and SS42-HA-dTMEM165-mCherry **(C)** generated using DeepTMHMM and represented as two-dimensional schematic models. Pink regions indicate epitopes predicted to be exposed to the cytosolic side of the membrane, whereas blue regions indicate regions located within the Golgi lumen. Predicted transmembrane domains are shown in red, and the signal peptide is indicated in orange. These in silico predictions provide a structural framework for interpreting experimental topology data. **(D)** Cells expressing the protein of interest are permeabilized using Triton X-100 (complete membrane permeabilization) or digitonin/SLO (selective permeabilization of the plasma membrane, preserving intracellular membranes such as the Golgi). Antibody accessibility is then assessed using antibodies directed either against endogenous epitopes of the protein or against engineered tags. In the absence of signal under Triton X-100 conditions, results are considered non-interpretable due to potential technical failure or lack of protein expression, and this condition serves as a control for assay validity. Under selective permeabilization, only cytosolic epitopes are accessible, whereas luminal or intramembrane epitopes are revealed only after Triton X-100 treatment. This differential accessibility allows determination of protein topology within cellular membranes.

**Figure S6.**
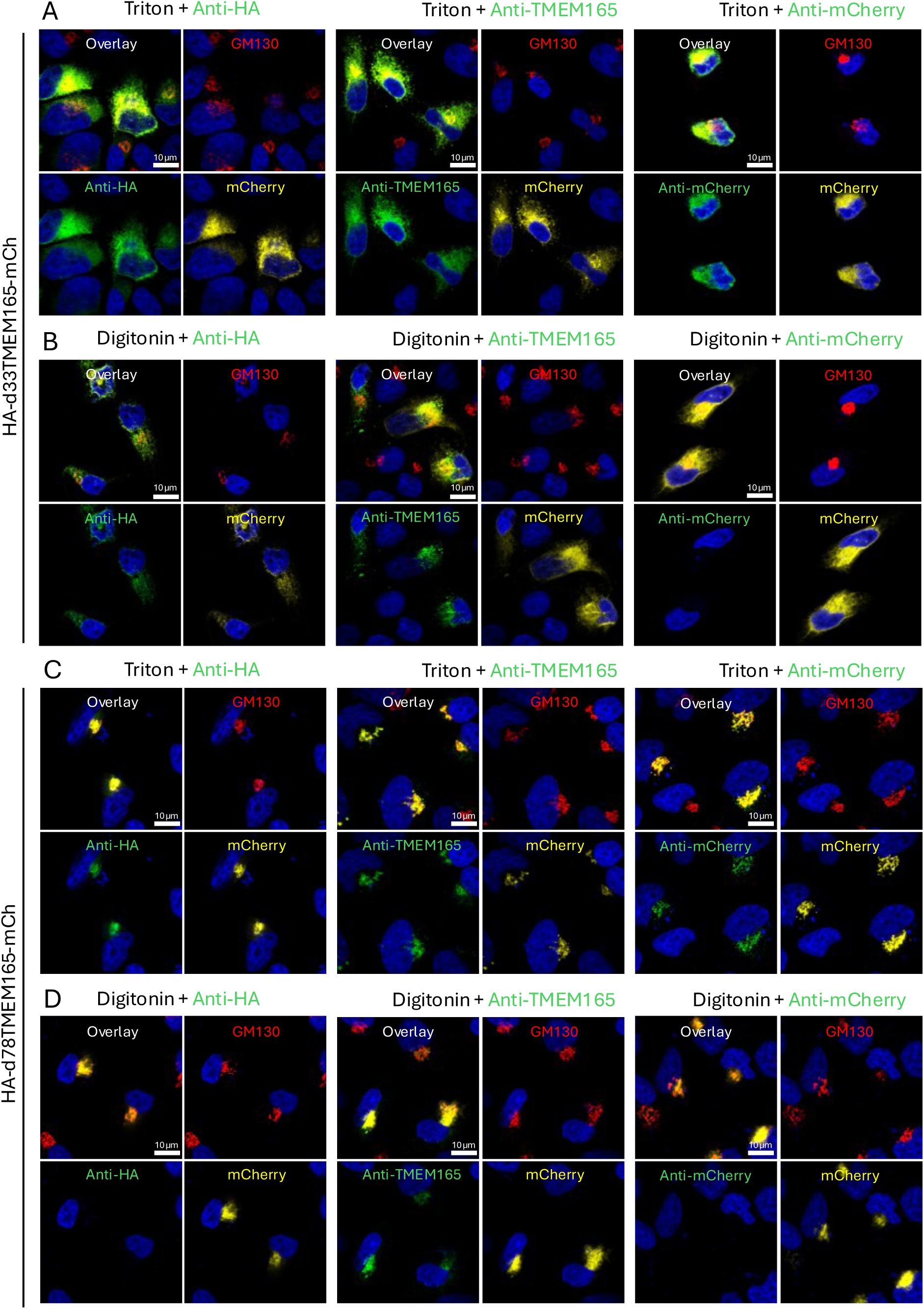
Representative fluorescence microscopy analysis of HA-Δ33TMEM165-mCherry and HA-Δ78TMEM165-mCherry constructs following Triton X-100 or digitonin permeabilization. **(A–D)** Representative fluorescence microscopy images of HA-Δ33TMEM165-mCherry **(A–B)** and HA-Δ78TMEM165-mCherry **(C–D)** constructs following Triton X-100 (A, C) or digitonin (B, D) permeabilization. For each panel, **left** correspond to staining with anti-HA, **middle** anti-TMEM165, and **right** anti-mCherry antibodies. GM130 staining (red) was used as a control for successful permeabilization of the plasma membrane, while intrinsic mCherry fluorescence (yellow) identified transfected cells. Green fluorescence corresponds to antibody staining. Only cells positive for both GM130 and mCherry signals were included in the analysis. Nucleus are highlighted with DAPI (blue).

**Figure S7.**
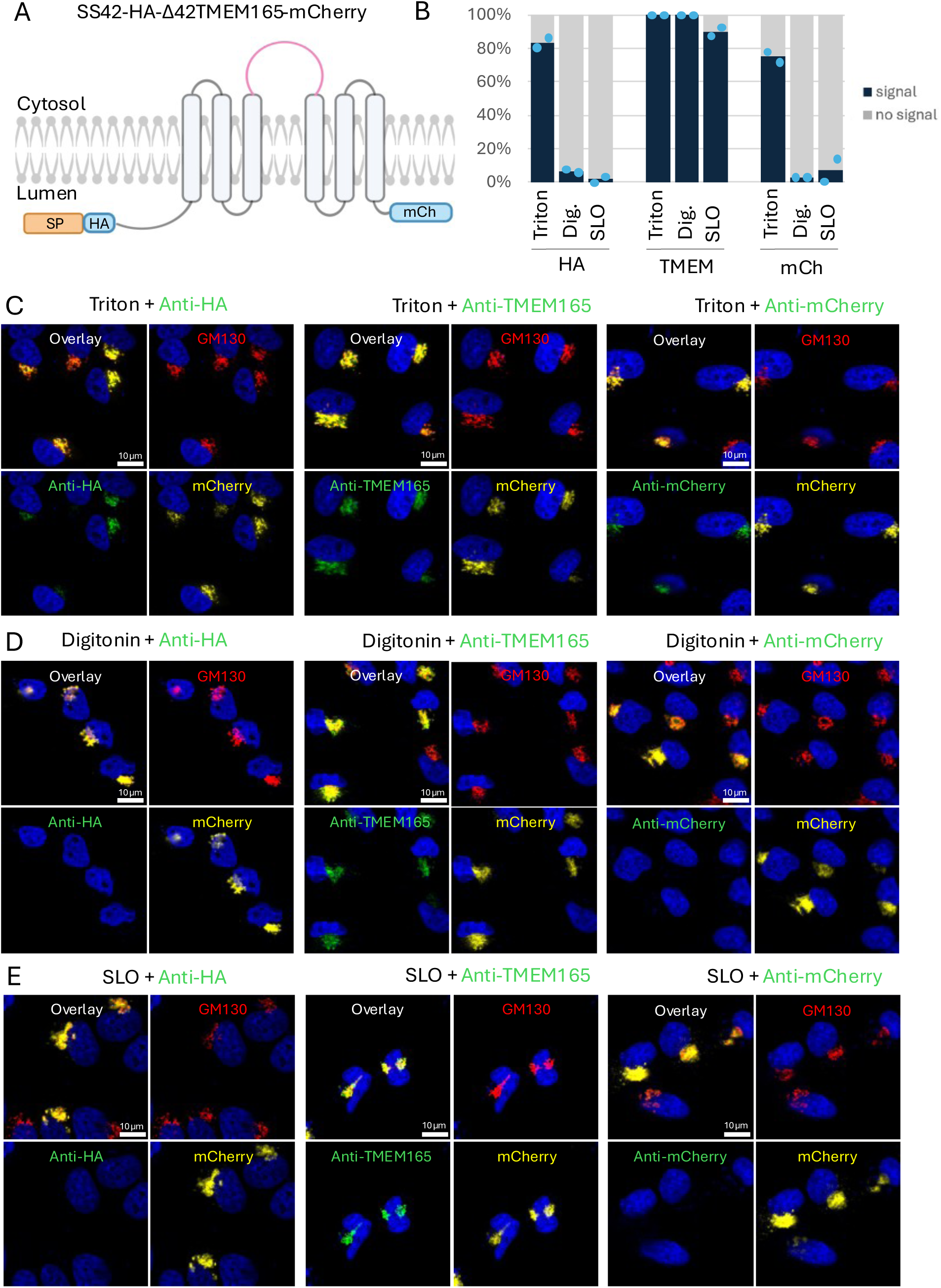
Membrane permeabilization analysis of the SS42-HA-d42TMEM165-mCherry construct using Triton X-100, digitonin, and streptolysin O (SLO). **(A)** Two-dimensional schematic representation of the predicted topology of the SS42-HA-d42TMEM165-mCherry construct within the Golgi membrane. Pink regions indicate epitopes exposed to the cytosolic side of the membrane, whereas blue regions indicate epitopes located within the Golgi lumen. Three epitopes were analyzed using specific antibodies: anti-HA recognizing the N-terminal HA tag, anti-TMEM165 recognizing a cytosolic loop of TMEM165, and anti-mCherry recognizing the C-terminal mCherry tag. **(B)** Percentage of cells displaying antibody staining (“signal”, dark blue) or no detectable staining (“no signal”, gray) following permeabilization with Triton X-100, digitonin, or SLO. Bars indicate the mean percentage obtained from two independent replicates, with a minimum of 50 cells analyzed per replicate. **(C–E)** Representative fluorescence microscopy images obtained following permeabilization with Triton X-100 **(C)**, digitonin **(D)**, or SLO **(E)**. For each permeabilization condition, **left** correspond to staining with anti-HA, **middle** anti-TMEM165, or **right** anti-mCherry antibodies. GM130 staining (red) was used as a control for successful plasma membrane permeabilization, while intrinsic mCherry fluorescence (yellow) allowed identification of transfected cells. Only cells positive for both GM130 and mCherry signals were included in the analysis. The presence or absence of the green signal was used to classify cells into the “signal” or “no signal” categories. Nucleus are highlighted with DAPI (blue).

---

**Table S1.**
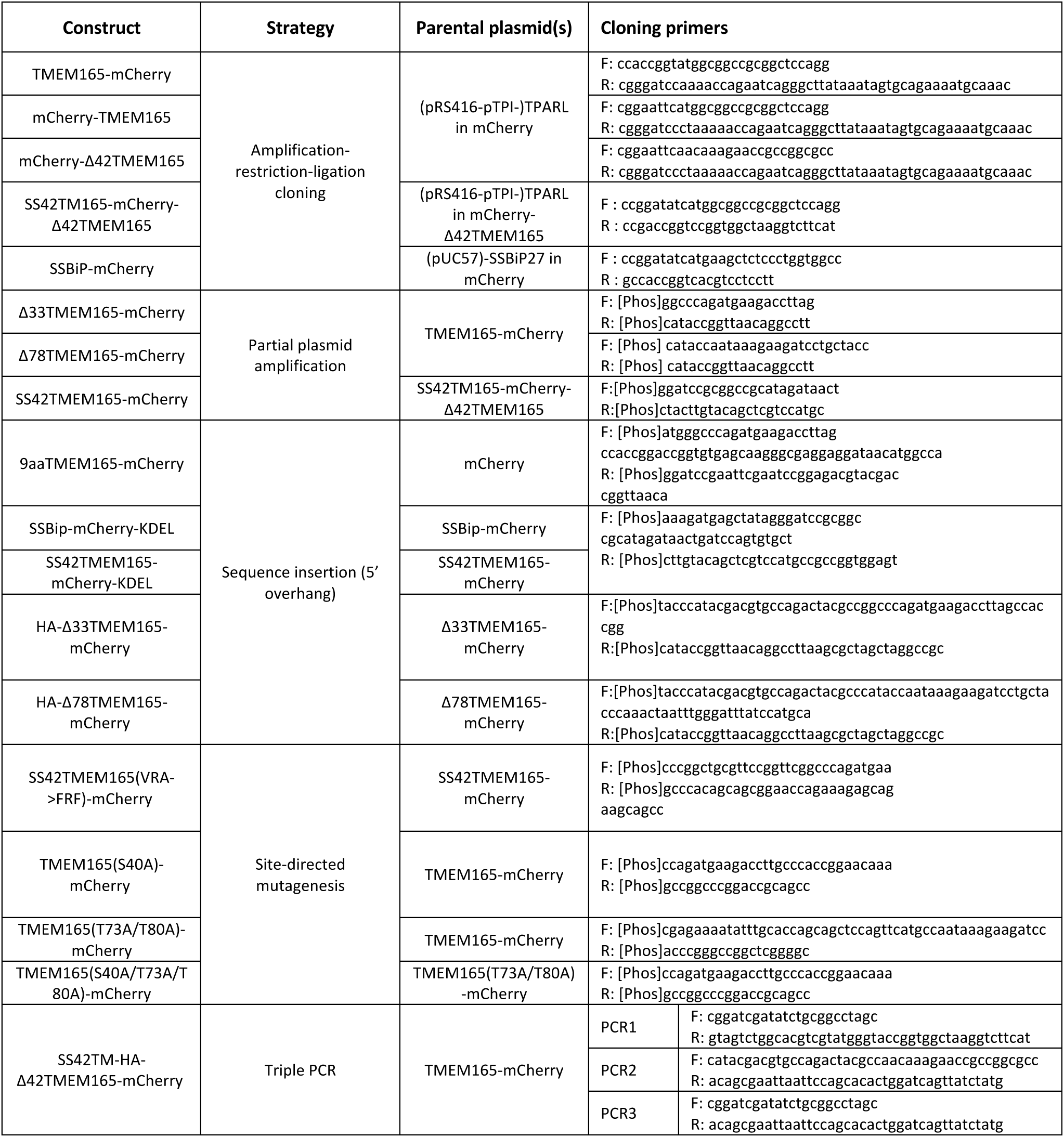
Cloning strategies, parental plasmids, and primers used for construct generation.

